# m6A modifications regulate intestinal immunity and rotavirus infection

**DOI:** 10.1101/2021.09.17.460776

**Authors:** Anmin Wang, Wanyin Tao, Jiyu Tong, Juanzi Gao, Jinghao Wang, Gaopeng Hou, Cheng Qian, Guorong Zhang, Runzhi Li, Decai Wang, Xingxing Ren, Kaiguang Zhang, Siyuan Ding, Wen Pan, Hua-Bing Li, Richard Flavell, Shu Zhu

## Abstract

N6-methyladenosine (m6A) is an abundant mRNA modification and affects many biological processes. However, how m6A levels are regulated during physiological or pathological processes such as virus infections, and the *in vivo* function of m6A in the intestinal immune defense against virus infections are largely unknown. Here, we uncover a novel antiviral function of m6A modification during rotavirus (RV) infection in small bowel intestinal epithelial cells (IECs). We found that rotavirus infection induced global m6A modifications on mRNA transcripts by down-regulating the m6a eraser ALKBH5. Mice lacking the m^6^A writer enzymes METTL3 in IECs (*Mettl3*^ΔIEC^) were resistant to RV infection and showed increased expression of interferons (IFNs) and IFN-stimulated genes (ISGs). Using RNA-sequencing and m6A RNA immuno-precipitation (RIP)-sequencing, we identified IRF7, a master regulator of IFN responses, as one of the primary m6A targets during virus infection. In the absence of METTL3, IECs showed increased *Irf7* mRNA stability and enhanced type I and III IFN expression. Deficiency in IRF7 attenuated the elevated expression of IFNs and ISGs and restored susceptibility to RV infection in *Mettl3*^ΔIEC^ mice. Moreover, the global m6A modification on mRNA transcripts declined with age in mice, with a significant drop from 2 weeks to 3 weeks post birth, which likely has broad implications for the development of intestinal immune system against enteric viruses early in life. Collectively, we demonstrated a novel host m6A-IRF7-IFN antiviral signaling cascade that restricts rotavirus infection *in vivo*.

## Introduction

N6-methyladenosine (m6A) is the most abundant internal mRNA modification and modulates diverse cellular functions through m6A-related writers, erasers, and readers[1-3]. The m6A modification directly recruits m6A-specific proteins of the YT521-B homology (YTH) domain family[1]. These proteins mediate the m6A-dependent regulation of pre-mRNA processing, microRNA processing, translation initiation, and mRNA decay[1]. In recent works, m6A modifications has been identified in the genomes of RNA viruses and the transcripts of DNA viruses with either a pro-viral or anti-viral role[4-8]. Furthermore, m6A RNA modification-mediated down-regulation of the α-ketoglutarate dehydrogenase (KGDH)-itaconate pathway inhibits viral replication independent of the innate immune response[9]. According to Gao et al. (2020), m6A modification preserves the self-recognition of endogenous transcripts. Deletion of the m6A writer *Mettl3* decreases the m6A modifications in endogenous retrovirus (ERV) transcripts. The accumulation of ERVs activates pattern recognition receptors (e.g. RIG-I) pathways, resulting in a detrimental interferon response in livers of fetal mice[10].

The m6A modification of the enterovirus 71 (EV71) RNA genome is important for viral propagation, and EV71 infection increases the expression of m6A writers *in vitro*[11]. m6A methyltransferase *Mettl3* knockdown reduces whereas m6A demethylase FTO knockdown increases EV71 replication[11]. In addition, human cytomegalovirus can up-regulate the expression of m6A-related proteins[12, 13]. Despite the knowledges about m6A regulation and function during viral infection revealed by these *in vitro* studies, the regulation of m6A modifications and the specific role of m6A in the anti-viral response *in vivo*, especially in the gastrointestinal tract, remains unclear.

Rotavirus (RV), a member of the family *Reoviridae*, is a nonenveloped icosahedral-structured virus with 11 segments of double-stranded RNA. Children under the age of five are at high risk of rotavirus infection, which causes severe diarrhea, dehydration, and death[14]. Rotaviruses encode multiple viral proteins to inhibit innate immune responses by degrading interferon regulatory factors (IRFs) and mitochondrial antiviral-signaling protein (MAVS), thus facilitating efficient virus infection and replication[15, 16]. The timely induction of an IFN response is key to the host successful control of invading viruses, including RV[17-19]. Here, we found rotavirus infection induced global m6A modifications on mRNA transcripts by down-regulating the m6a eraser ALKBH5. Mice lacking the m^6^A writer enzymes METTL3 in IECs (*Mettl3*^ΔIEC^) were resistant to RV infection. We identified IRF7, a master regulator of IFN responses[17], as one of the primary m6A targets during virus infection. In the absence of METTL3, IECs showed increased *Irf7* mRNA stability and enhanced type I and III IFN expression. Deficiency in IRF7 attenuated the elevated expression of IFNs and ISGs and restored susceptibility to RV infection in *Mettl3*^ΔIEC^ mice. Collectively, we identified a novel regulation and function of m6A modifications in an enteric viral infection model *in vivo*.

## Results

### The regulation and function of mRNA m6A modifications during rotavirus infection

Rotavirus infections primarily take place in children under the age of five in humans and in neonatal mice younger than 2 weeks-old[14, 20]. Intriguingly, total RNA m6A modifications in the mouse ileal tissues, revealed by a m6A dot blot, significantly declined from 2 weeks to 3 weeks post birth (Fig. 1a and 1b), which caused by increased *Alkbh5* expression (Fig. 1c). Besides, the global m6A RNA modification levels increased in the ileum tissue of suckling mice post RV murine strain EW infection (Fig. 1d and 1e). As a control, the global m6A levels decreased in *Mettl3* depleted bone marrow derived macrophages (BMDMs) (Fig. 1d). Thus, we hypothesize that RV may induce an enriched cellular m6A modification environment and a weakened innate immune response to facilitate virus replication. To investigate the *in vivo* role of m6A in the anti-RV immunity, we conditionally knocked out the m6A writer *Mettl3* in IECs (*Mettl3* f/f vil-cre, *Mettl3*^ΔIEC^), which is specifically infected by RV. Following infection with RV EW strain, the viral RNA load in *Mettl3*^ΔIEC^ mice ileum tissue was significantly lower than that in the wild-type (WT) littermates (Fig. 1f and s1). Fecal virus shedding was also significantly lower in *Mettl3*^ΔIEC^ mice (Fig. 1g). Genetic knockdown of *METTL3* in HT-29 cells, a human colonic epithelial cell line, by CRISPR-mediated gene silencing, also led to reduced RV replication (Fig. 1h), further highlighting the resistance phenotype to RV infection by METTL3 deficiency.

**Figure 1.**
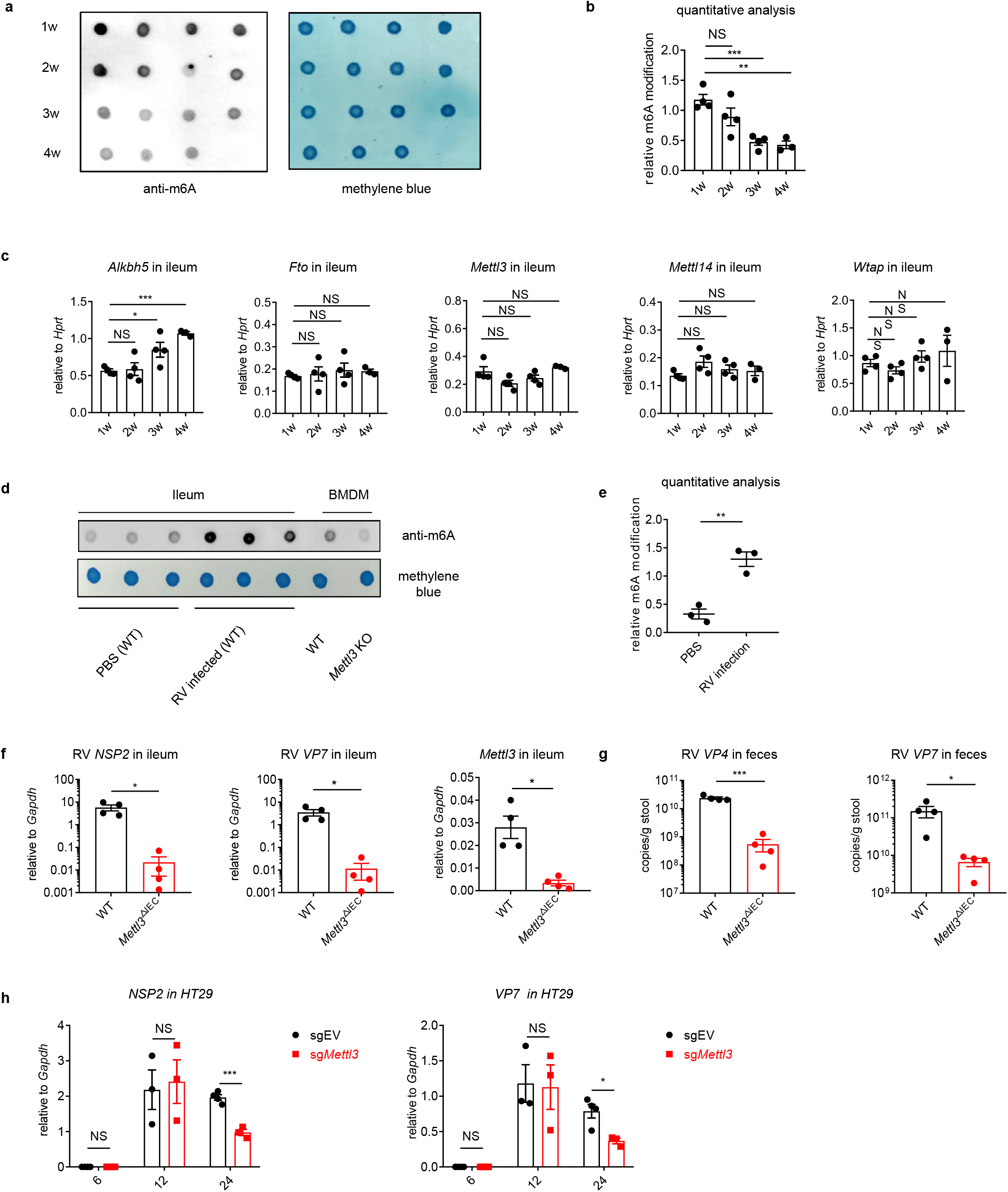
Rotavirus infection increases global m6A modifications, and *Mettl3* deficiency in intestinal epithelial cells results in increased resistance to rotavirus infection. (**a**) m6A dot blot analysis of total RNA in ileum tissues from mice with different ages. Methylene blue (MB) staining was the loading control. (**b**) Quantitative analysis of (a) (mean ± SEM), statistical significance was determined by Student’s t-test (**P < 0.005, ***P < 0.001, NS., not significant). The quantitative m6A signals were normalized to quantitative MB staining signals. (**c**) qPCR analysis of indicated genes in ileum tissues from mice with different ages (mean ± SEM). Statistical significance was determined by Student’s t-tests between groups (*P < 0.05, ***P<0.001, NS., not significant). (**d**) WT mice were infected by rotavirus EW strain at 8 days post birth. m6A dot blot analysis of total RNA in ileum tissue at 2 dpi. BMDMs from Wild type (WT) and *Mettl3* KO were used as the positive and negative controls, respectively. Methylene blue (MB) staining was the loading control. (**e**) Quantitative analysis of (c) (mean ± SEM). Statistical significance was determined by Student’s t-test (**P < 0.005). The quantitative m6A signals were normalized to quantitative MB staining signals. (**f-g**) 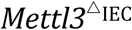 mice and littermate controls were infected by rotavirus EW strain at 8 days post birth. qPCR analysis of RV viral loads in ileum tissue (f) or fecal samples (g) from 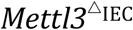 mice and littermate controls at 4 days post infection (dpi) (littermate WT n=4, 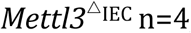, mean ± SEM). Statistical significance was determined by Student’s t-tests between genotypes (*P < 0.05, ***P < 0.001). (**h**) qPCR analysis of indicated genes in Rhesus rotavirus (RRV)-infected HT-29 cells transduced with *Mettl3* sgRNA or control sgRNA, at indicated hours post infection (hpi) (mean ± SEM), statistical significance was determined by Student’s t-test (*P < 0.05, ***P < 0.001, NS., not significant). Experiments in (**a-e**, and **h**) are repeated twice, (**f** and **g**) are repeated four times.

### *Mettl3* deficiency in IECs results in decreased m6A deposition on *Irf7*, and increased interferon responses

To dissect the underlying mechanism, we performed RNA-sequencing using the IECs from *Mettl3*-deficient mice and littermate controls at steady state. Most of the differentially expressed genes in *Mettl3*-deficient IECs vs WT IECs were enriched in the pathways of “defense response to virus”, “response to interferon-beta”, and “positive regulation of innate immune response” by gene ontology analysis (Fig. 2a). Heatmap also showed that a panel of interferon stimulate genes (ISGs) are upregulated in *Mettl3*-deficient IECs compared to WT IECs (Fig. 2b). To map potential m6A modification sites on mRNAs of these differential expressed genes in IECs, we conducted m6A RIP-sequencing based on a previously reported protocol [5]. We found that m6A modified one of the master regulators of IFNs, *Irf7* (Fig.2c), which played a key role in the network of these differential expressed genes in *Mettl3*-deficient IECs compared to WT IECs, analyzed by STRING (Fig.2d). Of note, IRF7 was the only IRFs that highly expressed in *Mettl3*-deficient IECs, and IRF7 was the prominently highest expressed IRFs in IECs (Fig. 2e), indicating IRF7 might be one of the key genes that are modulated by m6A modifications. In addition, the m6A peak was primarily located on the 5’ UTR and 3’ UTR of *Irf7* in ileum IECs (Fig. 2d). We also validated our results by m6A RIP-qPCR to examine m6A modification sites in *Irf7* mRNA based on our RIP-sequencing data and predicted results from the database (http://rna.sysy.edu.cn) (Fig. s2a and s2b). It should be noted that our m6A-RIP-seq did not identified previous reported IFN[12, 13], possibly due to the low expression level of *IFNb* in IECs at steady state.

**Figure 2.**
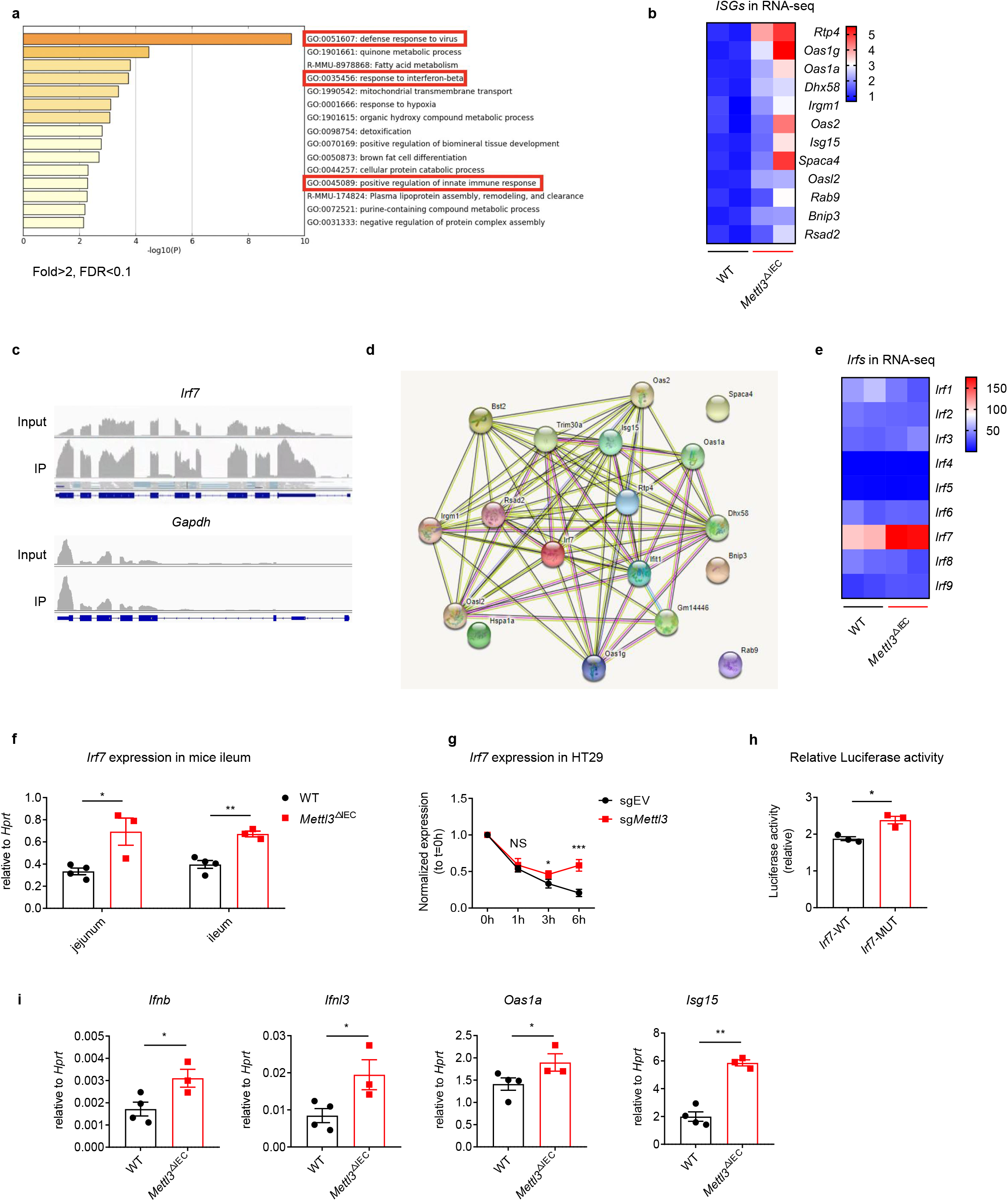
*Mettl3* deficiency in intestinal epithelial cells results in decreased m6A deposition on *Irf7*, and increased interferon responses. (**a**) Gene ontology (GO) analysis of differentially expressed genes in IECs from 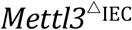 mice vs IECs from littermate WT mice. (**b**) Heat map of a subset of up-regulated *ISGs* in IECs from 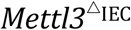 mice vs IECs from littermate WT mice, as revealed by RNA-seq (normalized data). (**c**) m6A-RIP-seq analysis of *Irf7* and *Gapdh* mRNA in the ileum of WT mice. (**d**) Gene regulation network of a subset of up-regulated genes including IRF7. (**e**) Heat map of Interferon regulatory factors (*Irfs*) in IECs from 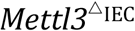 mice vs IECs from littermate WT mice, as revealed by RNA-seq (RPKM). (**f**) 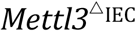mice and littermate controls were infected by EW at 8 days post birth. qPCR analysis of the *Irf7* expression in ileum and jejunum from 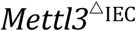 mice and littermate control at 2 dpi (littermate WT n=4, 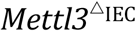 mice n=3, mean ± SEM). Statistical significance was determined by Student’s t-test (*P < 0.05, **P < 0.005). (**g**) q-PCR analysis of *Irf7* mRNA in *Mettl3* knockdown HT-29 cells or control cells in indicated time points post actinomycin D treatment (n=3, mean ± SEM). Statistical significance was determined by Student’s t-test (*P < 0.05, ***P < 0.001, NS., not significant). (**h**) Relative luciferase activity of HEK293T cells transfected with pmirGLO-*Irf7*-3’UTR (*Irf7*-WT) or pmirGLO-*Irf7*-3’UTR containing mutated m6A modification sites (*Irf7*-MUT). The firefly luciferase activity was normalized to Renilla luciferase activity. Statistical significance was determined by Student’s t-test (*P < 0.05). (**i**) 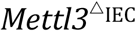 mice and littermate control and were infected by EW at 8 days post birth. qPCR analysis of selected *IFNs* and *ISGs* in ileum tissue at 2 dpi (littermate WT, n=4, 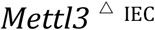 mice, n=3, mean ± SEM). Statistical significance was determined by Student’s t-tests between genotypes (*P < 0.05, **P < 0.005, ***P < 0.001). Experiments in (**f** and **i**) are repeated three times, (**g** and **h**) are repeated twice.

IRF7 is a known master regulator of Type I interferon and Type III interferon-dependent immune responses in respond to virus infection[16, 17, 21]. We reasoned that loss of m6A modification on Irf7 mRNA is responsible for the increased IFN response and subsequent resistance to RV infection. Thus, we first validated the regulation of *Irf7* mRNA levels by m6A in mice and in cells. We found an increase of *Irf7* mRNA in ileum tissue of *Mettl3*^ΔIEC^ mice compared to that in WT littermate control mice (Fig. 2f). Consistently, the expression of *Irf7* mRNA was also higher in *METTL3* knockdown HT-29 and *METTL3* knockout rhesus monkey MA104 cells, suggesting that the regulation of *Irf7* expression by m6A is likely conserved across species (Fig. s3a, s3b, s4a, and s4b). Furthermore, genetic knockdown of *METTL3* in HT-29 also led to increased IFN responses (Fig. s3). As m6A is known to regulate the mRNA decay, we next sought to determine whether the stability of *Irf7* mRNA is regulated by m6A. We used actinomycin D to block the *de novo* RNA synthesis in HT-29 cells to assess the RNA degradation by *METTL3* knockdown. The *Irf7* mRNA degraded significantly slower in *Mettl3-*knockdown HT-29 cells than the control cell line (Fig. 2g, s3b).

To directly evaluate the role of m6A in modulating the stability of *Irf7* mRNA, luciferase reporter assays were conducted. In comparison with wild-type *Irf7*-3’UTR (*Irf7*-WT) constructs, the ectopically expressed constructs harboring m6A mutant *Irf7*-3’UTR (*Irf7*-MUT) showed significantly increased luciferase activity (Fig. 2h). These results suggest that the upregulation of *Irf7* mRNA level in *Mettl3*^ΔIEC^ mice is caused by the loss of m6A modification mediated mRNA decay. To evaluate the potential influence of m6A on IRF7 transcriptional targets, we also measured the expressions of IFNs and ISGs in rotavirus infected ileum tissue from *Mettl3*^ΔIEC^ mice and littermate WT mice. We found the transcriptional targets of *Irf7*, were all up-regulated in *Mettl3*^ΔIEC^ mice (Fig. 2i). Furthermore, we found that the mRNAs of *Irf*7 and its transcriptional targets ISGs increased in the ileum of the mice from 1 to 4 weeks, with a dramatic up-regulation from 2 to 3 weeks (Fig. s5), which was concomitant with the decrease of global m6A modifications (Fig. 1a). These results demonstrated that METTL3 deficiency in IECs results in decreased m6A deposition on IRF7, and increased interferon response.

### IRF7 Deficiency attenuated the increased interferon response and resistance to rotavirus infection in *Mettl3*^ΔIEC^ mice

To determine whether *Irf7* plays a key role in the resistance phenotype to RV infection in *Mettl3* deficient mice in IECs, we crossed *Irf7*^−/−^ mice to *Mettl3*^ΔIEC^ mice. Following RV oral gavage, the expression of IFNs and ISGs in ileum from *Irf7*^−/−^ *Mettl3*^ΔIEC^ mice were significantly lower than those from *Mettl3*^ΔIEC^ mice at 2dpi (Fig 3a-c), and unlike the increased expression of IFNs and ISGs in *Mettl3*^ΔIEC^ mice vs littermate WT controls, by deficiency of IRF7, the expression of IFNs and ISGs in ileum from *Irf7*^−/−^ *Mettl3*^ΔIEC^ mice were not significantly different from those from *Irf7*^−/−^ mice (Fig 3a-c), suggesting that *Irf7* mediates the increased expression of IFNs and ISGs in *Mettl3*^ΔIEC^ mice.

**Figure 3.**
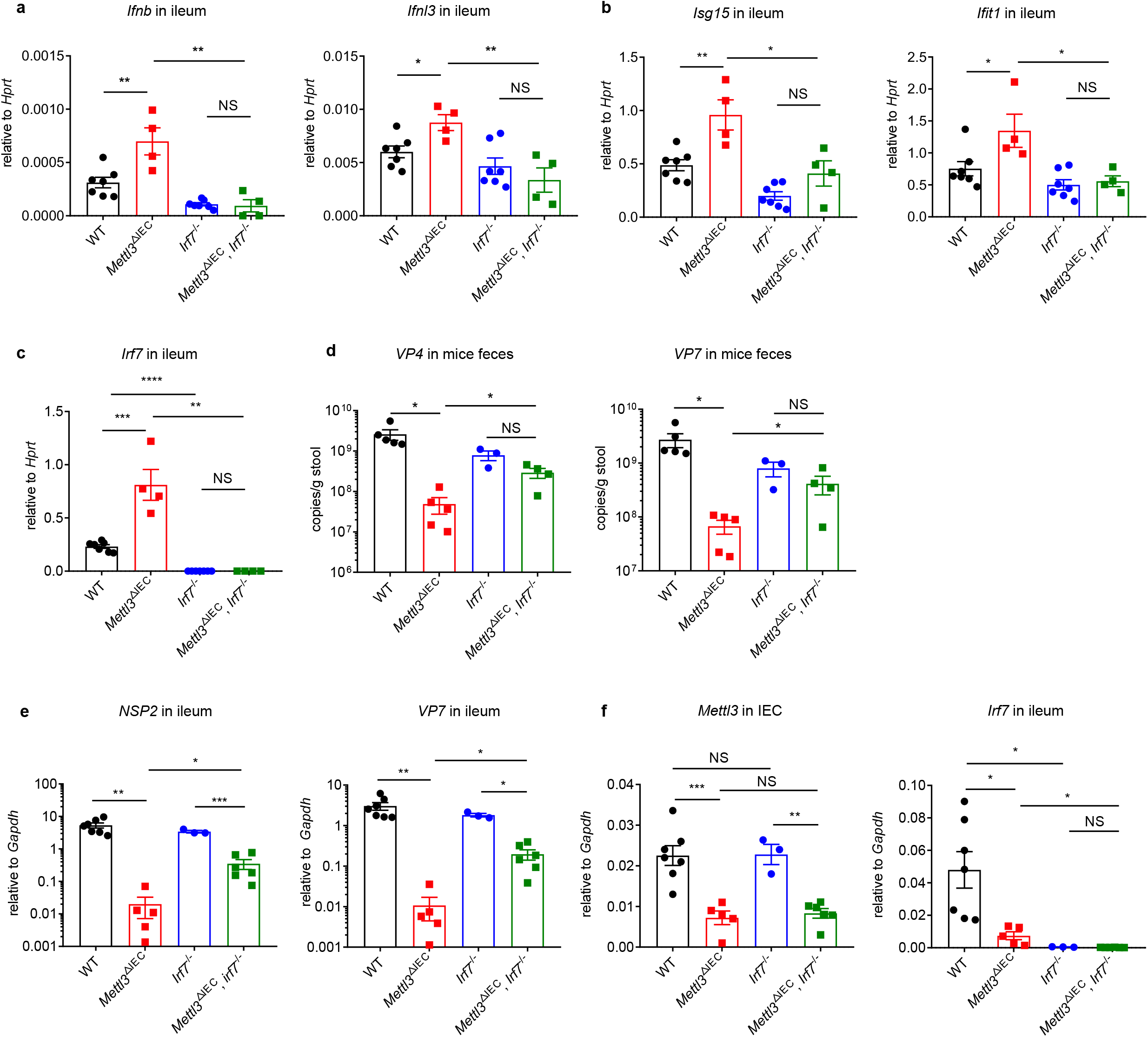
IRF7 Deficiency attenuated the increased interferon response and resistance to rotavirus infection in *Mettl3*^ΔIEC^ mice. (**a**-**c**) WT control mice, 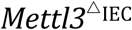 mice, *Irf7*^-/-^ mice and *Mettl3*^ΔIEC^*Irf7*^-/-^ mice are all littermates. They were infected by RV EW at 8 days post birth. qPCR analysis of selected *IFNs* (a), *ISGs* (b), or *Irf7* (c) expression in ileum from indicated groups of mice at 2 dpi (littermate WT n=7, *Mettl3*^ΔIEC^ n=5, *Irf7*^-/-^ n=3, *Mettl3*^ΔIEC^*Irf7*^-/-^ n=6, mean ± SEM). Statistical significance was determined by Student’s t-tests between genotypes (*P < 0.05, **P < 0.005, ***P < 0.001, ****P < 0.0001, NS., not significant). (**d**) qPCR analysis of fecal rotaviral shedding in indicated groups of mice at 4 dpi (littermate WT n=5, *Mettl3*^ΔIEC^ n=5, *Irf7*^-/-^ n=3, *Mettl3*^ΔIEC^*Irf7*^-/-^ n=4, mean ± SEM). Statistical significance was determined by Student’s t-tests between genotypes (*P < 0.05, NS., not significant). (**e-f**) qPCR analysis of RV proteins expression (e) or *Mettl3* and *Irf7* (f) in ileum from indicated groups of mice at 4 dpi (littermate WT n=7, *Mettl3*^ΔIEC^ n=5, *Irf7*^-/-^ n=3, *Mettl3*^ΔIEC^*Irf7*^-/-^ n=6, mean ± SEM). Statistical significance was determined by Student’s t-tests between genotypes and (*P < 0.05, **P < 0.005, ***P < 0.001, NS., not significant). Experiments in (**a-f**) are repeated twice.

Moreover, the *Irf7*^−/−^ *Mettl3*^ΔIEC^ mice showed significantly higher viral loads in ileum tissue and higher fecal shedding of RV *Mettl3*^ΔIEC^ (Fig. 3d-f). Similarly, unlike the much lower fecal viral shedding in *Mettl3*^ΔIEC^ mice vs littermate WT controls, by deficiency of IRF7, the fecal viral shedding in *Irf7*^−/−^ *Mettl3*^ΔIEC^ mice were not significantly different from those in *Irf7*^−/−^ mice (Fig 3d), suggesting that *Irf7* mediates the resistant phenotype of rotavirus infection measured by fecal viral shedding in *Mettl3*^ΔIEC^ mice. Notably, the viral proteins expression difference in ileum from *Irf7*^−/−^ *Mettl3*^ΔIEC^ mice vs that from *Irf7*^−/−^ mice (9.7-fold lower for NSP2 and 9.3-fold lower for VP7), was much lower than the viral proteins expression difference in ileum from *Mettl3*^ΔIEC^ mice vs that from littermate mice (267.1-fold lower for NSP2 and 283.4-fold lower for VP7) (Fig 3e), suggesting that besides the contribution of *Irf7* to the resistant phenotype of rotavirus infection in IECs from *Mettl3*^ΔIEC^ mice, other pathways (e.g. m6A modifications in RV RNA) may also play roles. Therefore, IRF7 is an important mediator of the increased IFNs and ISGs expression in *Mettl3*^ΔIEC^ mice and genetic deletion of *Irf7* restored the resistant phenotype of *Mettl3*^ΔIEC^ mice to RV infection.

### Rotavirus suppresses ALKBH5 expression through NSP1 to evade immune defense

We next sought to determine how RV regulates the m6A modifications in IECs. We first measured whether RV infection regulates the m6A-related writer and eraser proteins in the intestine. The protein levels of the methyltransferases METTL3 and METTL14 and demethylase FTO were not affected by RV infection in ileum tissue (Fig. 4a and b). In contrast, the protein level of demethylase ALKBH5 was significantly down-regulated by RV infection in the ileum (Fig. 4a, 4b). To determine whether ALKBH5 play a role in anti-RV infection since it’s suppressed during RV infection, we generated the IEC-specific deletion of ALKBH5 in mice (*Alkbh5* f/f vil-cre, *Alkbh5*^ΔIEC^). The depletion of ALKBH5 in IECs did not affected the anti-RV immune response (Fig. 4c), the viral shedding in the feces (Fig.4d), or the viral protein expression in the ileum (Fig. 4e), likely due to the suppressed ALKBH5 expression in ileum tissue of WT mice infected by RV.

**Figure 4.**
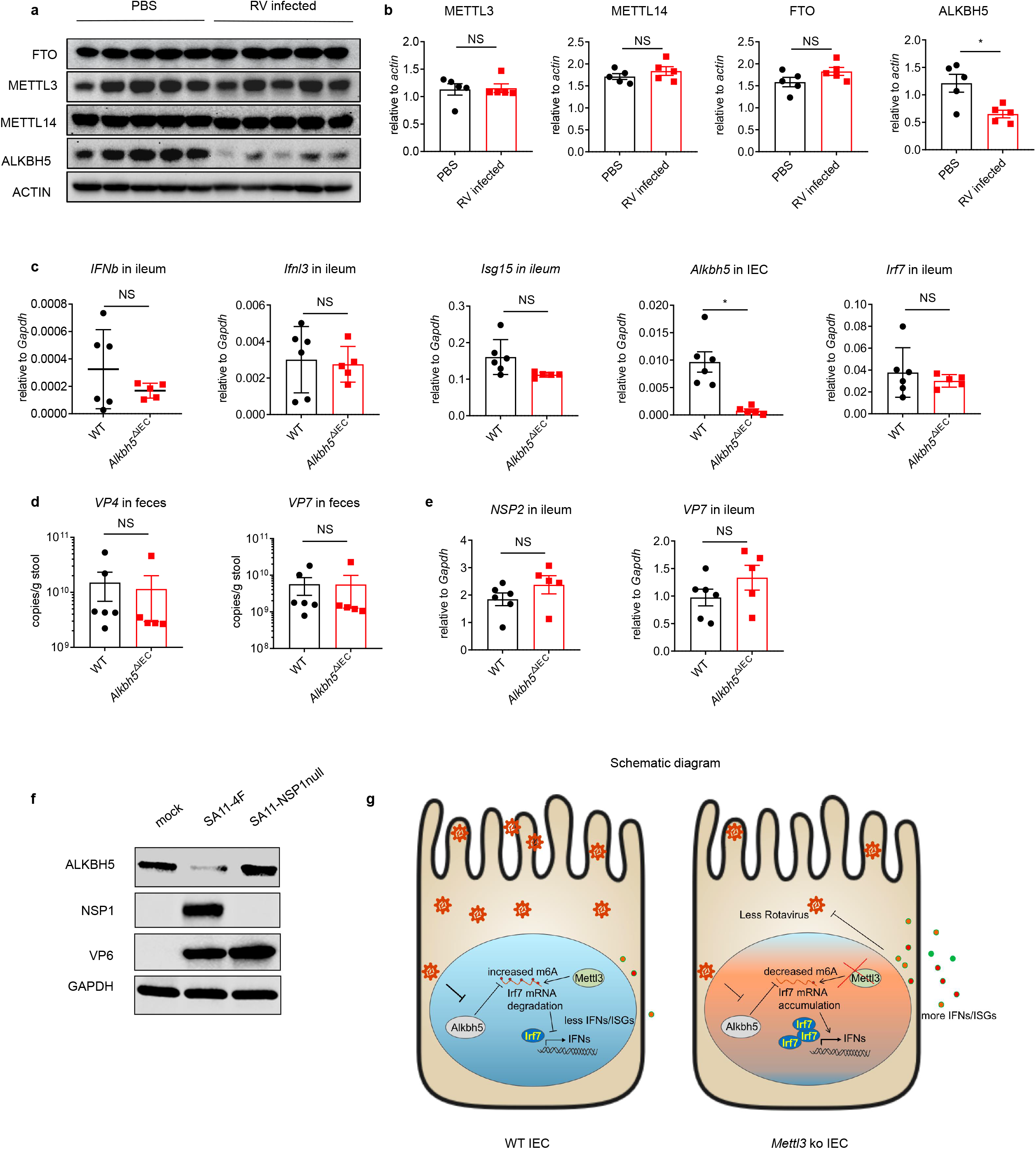
Rotavirus suppresses ALKBH5 expression through NSP1 to evade immune defense. (**a**) WT mice were infected by RV EW at 8 days post birth. Immunoblotting with antibodies target ALKBH5, FTO, METTL14 and METTL3 in ileum tissue from mice infected with RV EW at 2 dpi or treated with PBS. (**b**) Quantitative analysis of (a)(mean ± SEM), Statistical significance was determined by Student’s t-test (*P < 0.05, NS., not significant). (**c-e**) 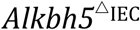 mice and littermate controls were infected by RV EW at 8 days post birth. qPCR analysis of indicated genes expression in ileum (c), viral shedding in feces (d), and viral proteins expression in ileum (e), from 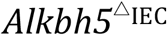 mice or littermate controls at 4 days post infection (littermate WT n=6, 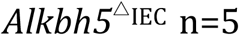, mean ± SEM). Statistical significance was determined by Student’s t-tests between genotypes (*P < 0.05, NS., not significant). (**f**) Immunoblotting with antibodies target ALKBH5, NSP1, VP6 and GAPDH in HEK293 cells infected by SA11-4F and SA11-NSP1null (MOI=1) for 24h. (**g**) Graphical abstract illustrating the functions and molecular mechanisms of m6A modifications on *Irf7* in anti-RV infection. Experiments in (**a**-**e**) are repeated three time, (**f**) are repeated twice.

Non-structural protein 1 (NSP1) is a well-established RV-encoded innate immune antagonist that are showed to degrade IRF3 and β-Trcp [22, 23]. To test the potential role of NSP1 in AlKBH5 inhibition, we used the recently developed reverse genetics system and rescued recombinant WT RV SA11 strain and a mutant virus that does not express NSP1 (NSP1-null)[24]. We infected HEK293 cells with WT and NSP1-null RVs, only WT RV reduced ALKBH5 protein levels (Fig. 4f), suggesting that the down-regulation of ALKBH5 expression by RV is NSP1-dependent. These results suggest that RV might evades the anti-viral immune response via downregulation of ALKBH5 expression.

## Discussion

Previous studies reported that m6A modifications on mRNA in mice embryonic fibroblasts or normal human dermal fibroblasts negatively regulate the IFN response by accelerating the mRNA degradation of type I IFNs [12, 13]. However, these studies were mainly conducted in vitro, leaving the relationship between m6A and IFN pathway *in vivo* an unexplored territory. Since type I and type III IFNs play a critical role in the antiviral immune response in the GI tract[19], we used an RV infection model, in which IECs are specifically infected, as well as conditional knockout mice with IECs-specific depletion of m6A writer METTL3 or m6A eraser ALKBH5, to study the role of m6A modification in regulating IFN response towards rotavirus in IECs. Using RNA-seq and m6A-seq techniques, we identified IRF7, a key transcription factor upstream of IFNs and ISGs, as one of the m6A modified targets during RV infection, and IRF7 is essential to mediate the elevated anti-rotavirus immune response by METTL3 deficiency. These results identify *Irf7* as an important m6A target, and characterize, for the first time, the regulation of IFN response during RV infection in intestine, enabling a better understanding of how m6A modifications on mRNAs regulate anti-viral innate immune responses.

In addition to directly regulating target genes that involved in innate immune pathways, m6A modification is also known to affect viral gene expression, replication and generation of progeny virions[1, 25]. Some viral RNA genomes are modified by m6A, such as simian virus 40[4, 26], influenza A virus[27], adenovirus[28], avian sarcoma virus[29], Rous Sarcoma virus[30], hepatitis C virus[31], and Zika virus[32]. Through our m6A-RIP-qPCR, Rotavirus RNA was also found to have m6A modification (Figure s6). Of note, genetic deletion of METTL3 in monkey kidney MA104 cell line, which has limited IFN responses[33], also led to reduced RV replication. Further, IRF7 deficiency didn’t fully restore the suppressed rotaviral infection in *Mettl3*^ΔIEC^ mice (Fig 3e). These results suggest that besides the contribution of *Irf7* to the resistant phenotype of rotavirus infection in IECs from *Mettl3*^ΔIEC^ mice, other pathways (e.g. m6A modifications in RV RNA) may also play roles. The detailed mechanisms warrant further investigations in the future.

Intriguingly, m6A modifications were shown to maintain the genomic stability of mice embryonic stem cells by promoting the degradation of endogenous retrovirus (ERV) mRNA, or by regulating ERV heterochromatin and inhibiting its transcription[2, 3]. The absence of m6A results in the formation of abnormal endogenous dsRNA, which causes an aberrant immune response and necrosis in the hematopoietic system[10]. In the intestine, through immunostaining with J2 antibody, we detected the increase of dsRNA levels in IECs of *Mettl3*^ΔIEC^ mice compare to the littermate WT mice (Figure s7). In consistency, the aforementioned RNA-seq data showed that the expression of a set of ISGs including IRF7 was significantly upregulated in *Mettl3* KO IECs. Our data suggest a dual activation model in steady state that in the absence of *METTL3*, the increase of dsRNA will induce the IFN responses, and the increase of mRNA stability of *Irf7*, which serves as a key transcription factor of IFN and ISG expression, will amplify this process. Furthermore, m6A modification of viral RNA genomes affect the activation of innate sensor-mediated signaling[34, 35]. Decreased m6A modification on RV genomes may activate innate sensors directly and induce higher IFN response, which will also be amplified by increase of mRNA stability of *Irf7*.

Although m6A is involved in many important biological processes, the regulation of m6A modifications remain poorly understood. Here, we found RV infection down-regulates the level of m6A eraser ALKBH5 to induce m6A modifications, in an NSP1 dependent manner. The precise mechanism remains to be examined. As a result, *ALKBH5* deficiency in IECs results in normal susceptibility to RV infection. In addition, the global m6A modification on mRNA transcripts declines by ages in the intestine, with a significant drop from 2 to 3 weeks post birth, which implicate the drop of RV infectivity in adult mice vs neonatal mice. The dual regulation of m6A levels during RV infection and development provide new insights into the choice of either RV or the host on regulation of m6A modification to achieve either immune evasion or immune surveillance, respectively.

In conclusion, our work shed light into a novel role of m6A modifications in RV infection *in vivo*, and reported a tissue specific regulation of m6A during RV infection or development (Figure 4g). Future studies on tissue-specific regulation of m6A modification by viral infections in other tissues and organs (e.g. lung, liver) will be of interest.

## Materials and Methods

### Mice

Mettl3 conditional knockout mice were generated by inserting two loxp sites into the intron after the first exon and the intron before the last exon of *Mettl3* using CRISPR/cas9 based genome-editing system as previously described[5]. Alkbh5 conditional knockout mice were generated by inserting two loxp sites into the introns flanking the first exon of *Alkbh5* using CRISPR/cas9 based genome-editing system as previously described[36]. Genotyping of *Mettl3* f/f mice, *Alkbh5* f/f mice, Vil-cre mice (The Jackson Laboratory, Stock No: 021504), and *Irf7*^-/-^ mice (RIKEN BRC, RBRC01420) were confirmed by PCR using primers as list below:

*Mettl3* f/f mice

Mettl3-L1+:CCCAACAGAGAAACGGTGAG

Mettl3-L2-:GGGTTCAACTGTCCAGCATC

Vil-cre mice

Vil-Cre-182/150-F:GCCTTCTCCTCTAGGCTCGT

Vil-Cre-182-R:TATAGGGCAGAGCTGGAGGA

Vil-Cre-150-R:AGGCAAATTTTGGTGTACGG

*Irf7*^-/-^ mice

RBRC01420-Irf7-WT-F:GTGGTACCCAGTCCTGCCCTCTTTATAATCT

RBRC01420-Irf7-Mut-F:TCGTGCTTTACGGTATCGCCGCTCCCGATTC

RBRC01420-Irf7-R:AGTAGATCCAAGCTCCCGGCTAAGTTCGTAC

*Alkbh5* f/f mice

Alkbh5-L1+: GCACAGTGGAGCACATCATG

Alkbh5-L2-: CAGAGGGCAAGCAACCACAC

The sex-, age- and background-matched littermates of the knockout or conditional knockout mice were used as the controls in the present study. All mice were on the C57BL/6 background. Mice were maintained in SPF conditions under a strict 12 h light cycle (lights on at 08:00 and off at 20:00). All animal studies were performed according to approved protocols by the Ethics Committee at the University of Science and Technology of China (USTCACUC202101016).

### Cell culture

The MA104 cell line was obtained from the Cell Resource Center, Peking Union Medical College (which is the headquarter of National Infrastructure of Cell Line Resource). The identity of the cell line was authenticated with STR profiling (FBI, CODIS). The results can be viewed on the website (http://cellresource.cn). HEK293T (ATCC CRL-3216), HT-29 (ATCC HTB-38D™) were obtained from the American Type Culture Collection (ATCC). All of these cells were cultured in Dulbecco’s modified Eagle’s medium (DMEM) (Hyclone) supplemented with 10% fetal bovine serum (FBS) (Clark); All cells were cultured at 37°C in 5% CO2.

### Plasmids and SgRNAs

All gene silencing was done using a CRISPR–cas9 system, with lentiCRISPR v2 plasmid (Addgene no. 52961). The following sgRNAs were cloned downstream of the U6 promoter:

Human, rhesus and mouse METTL3: 5′ -GGACACGTGGAGCTCTATCC-3′;

Lentiviruses were generated by co-transfection of lentiCRISPR v2 constructs and packaging plasmids (psPAX2, Addgene no. 12260 and pMD2.G, Addgene no. 12259), using PEI DNA transfection reagent (Shanghai maokang biotechnology), into HEK293T cells, according to the manufacturer’s instructions. At 48h post transfection, supernatants were collected and filtered through a 0.22μm polyvinylidene fluoride filter (Millex). To induce gene silencing, cells were transduced with lentivirus expressing sgRNA and were puromycin selected (2μg/ml) for 4–5 days. The depletion of target proteins was confirmed by immunoblot analysis.

### Virus infections

Rhesus and simian RV strains, including RRV (Rhesus), SA11-4F (simian), SA11-NSP1null (simian) were propagated in MA104 cells as previously described[23]. Viruses were activated by trypsin (5 μg/ml) at 37°C for 30 min prior to infection. Cells were washed with PBS three times and incubated with RV at different MOIs at 37°C for 1 hr. After removal of RV inoculum, cells were washed with PBS, cultured in serum-free medium (SFM) and harvested for qPCR and western blot analysis.

EW stock virus was prepared by infecting 5-days-old C57BL/6J mice, and harvesting crude centrifugation-clarified intestinal homogenate as previously described[15].

For all rotavirus infection except indicated elsewhere, 8-day-old wild-type mice, or genetically deficient mice were orally inoculated by gavage with RV EW virus as previously described[15]. Mice were sacrificed, stool and small intestinal tissue were collected at indicated time points post infection. Viral loads in intestinal tissues and feces were detected by RT–qPCR.

### RT-qPCR

For cells and tissues, total RNA was extracted with TRNzol Universal reagent (Tiangen) in accordance with the manufacturer’s instructions. Real-time PCR was performed using SYBR® Premix Ex Taq™ II (Tli RNaseH Plus) (Takara) and complementary DNA was synthesized with a PrimeScript™ RT reagent Kit with gDNA Eraser (Takara). The target genes were normalized to the housekeeping gene (*Gapdh, HPRT*) shown as 2−ΔCt. The used primers are as follows:

Primers detect mouse genes:

Mettl3-F: ATTGAGAGACTGTCCCCTGG

Mettl3-R: AGCTTTGTAAGGAAGTGCGT

Mettl14-F: AGACGCCTTCATCTCTTTGG

Mettl14-R: AGCCTCTCGATTT CCTCTGT

Fto-F: CTGAGGAAGGAGTGGCATG

Fto-R: TCTCCACCTAAGACTTGTGC

Alkbh5-F: ACAAGATTAGATGCACCGCG

Alkbh5-R: TGTCCATTTCCAGGATCCGG

Wtap-F: GTTATGGCACGGGATGAGTT

Wtap-R: ATCTCCTGCTCTTTGGTTGC

Gapdh-F: TGAGGCCGGTGCTGAGTATGTCG

Gapdh-R: CCACAGTCTTCTGGGTGGCAGTG

Hprt-F: ACCTCTCGAAGTGTTGGATACAGG

Hprt-R: CTTGCGCTCATCTTAGGCTTTG

Irf7-F: GCTCCAGTGACTACAAGGCAT

Irf7-R: TTGGGAGTTGGGATTCTGAG

Isg15-F: GGTGTCCGTGACTAACTCCAT

Isg15-R: TGGAAAGGGTAAGACCGTCCT

Oas1a-F: GCCTGATCCCAGAATCTATGC

Oas1a-R: GAGCAACTCTAGGGCGTACTG

Ifnb-F: TCCGAGCAGAGATCTTCAGGAA

Ifnb-R: TGCAACCACCACTCATTCTGAG

Ifnl3-F: AGCTGCAGGCCTTCAAAAAG

Ifnl3-R: TGGGAGTGAATGTGGCTCAG

Primers detect human genes:

hGAPDH-F: ATGACATCAAGAAGGTGGTG

hGAPDH-R: CATACCAGGAAATGAGCTTG

hIRF7-F: CGAGACGAAACTTCCCGTCC

hIRF7-R: GCTGATCTCTCCAAGGAGCC

hIFNL3-F: TAAGAGGGCCAAAGATGCCTT

hIFNL3-R: CTGGTCCAAGACATCCCCC

hCXCL10-F: TGGCATTCAAGGAGTACCTC

hCXCL10-R: TTGTAGCAATGATCTCAACACG

hIFIT1-F: CAACCATGAGTACAAATGGTG

hIFIT1-R: CTCACATTTGCTTGGTTGTC

Primers detect rhesus genes:

Rhesus-GAPDH-F: ATGACATCAAGAAGGTGGTG

Rhesus-GAPDH-R: CATACCAGGAAATGAGCTTG

Rhesus-IFIT1-F: CAACCATGAGTACAAATGGTG

Rhesus-IFIT1-R: CTCACACTTGCTTGGTTGTC

Rhesus-IRF7-F: GTTCGGAGAGTGGCTCCTTG

Rhesus-IRF7-R: TCACCTCCTCTGCTGCTAGG

Rhesus-IFNL1-F: ACTCATACGGGACCTGACAT

Rhesus-IFNL1-R: GGATTCGGGGTGGGTTGAC

Rhesus-IFNb-F: GAGGAAATTAAGCAGCCGCAG

Rhesus-IFNb-R: ATTAGCAAGGAAGTTCTCCACA

Primers detect virus genes:

Rotavirus EW-NSP2-F: GAGAATGTTCAAGACGTACTCCA

Rotavirus EW-NSP2-R: CTGTCATGGTGGTTTCAATTTC

Rotavirus EW-VP4-F: TGGCAAAGTCAATGGCAACG

Rotavirus EW-VP4-R: CCGAGACACTGAGGAAGCTG

Rotavirus EW-VP7-F: TCAACCGGAGACATTTCTGA

Rotavirus EW-VP7-R: TTGCGATAACGTGTCTTTCC

RRV VP7-F: ACGGCAACATTTGAAGAAGTC

RRV VP7-R: TGCAAGTAGCAGTTGTAACATC

RRV NSP2-F: GAGAATCATCAGGACGTGCTT

RRV NSP2-R: CGGTGGCAGTTGTTTCAAT

RRV NSP5-F: CTGCTTCAAACGACCCACTCAC

RRV NSP5-R: TGAATCCATAGACACGCC

### m6A dot blot assay

Total RNA was isolated from mice ileum using TRNzol Universal Reagent (Tiangen, Lot#U8825) according to the manufacturer’s instructions. RNA samples were quantified using UV spectrophotometry and denatured at 65 °C for 5 min. The m6A-dot-blot was performed according to a published work[37]. In brief, the primary rabbit anti-m6A antibody (1:5000, Synaptic System, #202003) was applied to the Amersham Hybond-N+ membrane (GE Healthcare, USA) containing RNA samples. Dot blots were visualized by the imaging system after incubation with secondary antibody HRP-conjugated Goat anti-rabbit IgG (Beyotime, A0208).

### Western blot

Briefly, cells and tissue were lysed with RIPA buffer (Beyotime Biotechnology) supplemented with PMSF (Beyotime Biotechnology) and protease inhibitor cocktail (Roche). Mettl3 (abcam, ab195352, 1:2000), Mettl14 (sigma, HPA038002, 1:2000), ALKBH5 (sigma, HPA007196, 1:2000), FTO (abcam, ab92821), NSP1 and VP6 (gift from Harry B. Greenberg lab), Gapdh (proteintech), and beta-actin(proteintech) antibodies were used in accordance with the manufacturer’s instructions. After incubation with the primary antibody overnight, the blotted PVDF membranes (Immobilon, IPVH00010) were incubated with goat anti-rabbit IgG-HRP (Beyotime, A0208) or goat anti-mouse IgG-HRP (Beyotime, A0216) and exposed with BIO-RAD ChemiDocTM Imaging System for a proper exposure period.

### RNA degradation assay

The stability of targeted mRNA was assessed as previously described[5]. In brief, *Mettl3* knock down HT-29 and control cell were plated on 24-well plate. Actinomycin-D (MCE, HY17559) was added to a final concentration of 5μM, and cells were harvested by indicated time points after actinomycin-D treatment. The RNA samples are processed and qPCR was used to measure the mRNA transcripts, all data were normalized to that of t=0 time point.

### Dual-luciferase assay

pmirGLO (Firefly luciferase, hRluc) vector of the Dual-luciferase Reporter assay system (Promega, E1910) was used to determine the function of m6A modification within the 3’UTR of Irf 7 transcripts. The potential m6A modification sites were predicted on SRAMP website. The assay was performed according to the manufacture’s instruction: Briefly, 300 ng of pmirGLO vector containing Irf7-3’UTR or m6A-mutant Irf7-3’UTR were transfected into HEK293T cells in triplicate wells. The relative luciferase activity was accessed 36 h post-transfection.

### Isolation of IECs in the intestine

Small intestines were excised and flushed thoroughly three times with PBS. They were turned inside out and cut into ∼1 cm sections then transferred into RPMI with 2 mM EDTA, and shaken for 20 min at 37 °C. Supernatants were collected through a 100-mm cell strainer to get single-cell suspensions. Cells were collected as the IEC fraction which contains both epithelial cells (∼90%) and lymphocytes (IEL, ∼10%). Single-cell suspension was used for further analysis.

### RNA-Seq

IECs from *Mettl3*^ΔIEC^ mice as well as the wild-type littermate control mice were isolated as described in previous section. Total RNAs were extracted with TRNzol universal RNA Reagen kits. Berrygenomics (Beijing, China) processed the total RNA and constructed the mRNA libraries, and subject them to standard illumine sequencing on Novaseq 6000 system, and obtained > 40 million Pair-end 150 reads for each sample. Raw RNA-sequencing reads were aligned to the mouse genome (mm10, GRCm38) with STAR (v2.5.3a). Gene expression levels and differential analysis was performed with edgeR(v3.29.2). Genes were considered significantly differentially expressed if showing ≥1.5-fold change and FDR < 0.05. Gene set analysis was performed and enriched pathways were obtained through online bioinformatics tools (metascape) and GSEA (v4.0.3). Pathway plot were gene-rated with R package ‘ggplot2’[5].

### m6A RNA-IP-qPCR & m6A RNA-IP-Seq

m6A RNA-IP-Seq was carried out according to a previously published protocol[5]. In brief, total cellular RNA extracted from WT C57 mice IEC was fragmented by ZnCl2 followed by ethanol precipitation. Fragmented RNA was incubated with an anti-m6A antibody (Sigma Aldrich ABE572) and IgG IP Grade Rabbit polyclonal antibody (abcam, lot: 934197). The eluted RNA and input were subjected to high-throughput sequencing using standard protocols (Illumina, San Diego, CA, USA) or processed as described in ‘RT-qPCR’ section, except that the data were normalized to the input samples. The m6A RIP-Seq data were analyzed as described previously[5].

RIP-ptpn4-F: CCTCCCATCCCGGTCTCCACC

RIP-ptpn4-R: GGCTGCCCATCTTCAGGGGT

RIP-RPS14-F: ACCTGGAGCCCAGTCAGCCC

RIP-RPS14-R: CACAGACGGCGACCACGACG

m6A-IRF7-F: GACAGCAGCAGTCTCGGCTT

m6A-IRF7-R: ACCCAGGTCCATGAGGAAGT

m6A sites on RV-EW RNA were predicted on http://www.cuilab.cn/sramp website, and m6A-RIP-qPCR primer were designed on NCBI primer blast according to the predicted m6A sites.

m6A-RIP-qPCR primer:

RIP-EW-VP1-F: ACGAAATGCTTGTTGCTATGAGT

RIP-EW-VP1-R: AACCTGTCCGTCAACCATTC

RIP-EW-VP2-F: GGCCAGAACAGGCTAAACAAC

RIP-EW-VP2-R: CGCAGTTCTCTTTCGCCATTT

RIP-EW-VP3-F: CGATGACAGCACAAAAGTCGG

RIP-EW-VP3-R: CGTGTCTCTTGCGAAGTC

RIP-EW-VP4-F: TCAGCAGACGGTTGAGACTG

RIP-EW-VP4-R: GGCTGAGATGTCATCGAAGTT

RIP-EW-NSP1-F: CCTCACATCTCTGCTACATGAACT

RIP-EW-NSP1-R: TGCTGGTTGGACATGGAATGA

RIP-EW-VP6-F: CTGCACTTTTCCCAAATGCTCA

RIP-EW-VP6-R: GAGTCAATTCTAAGTGTCAGTCCG

RIP-EW-NSP3-F: CTTGACGTGGAGCAGCAAC

RIP-EW-NSP3-R: AATGTTTCAATGTCGTCCAACG

RIP-EW-NSP2-F: TCCACCACTCTAAAGAACTACTGC

RIP-EW-NSP2-R: TCCGCTGTCATGGTGGTTTC

RIP-EW-VP7-F: TCGGAACTTGCAGACTTGAT

RIP-EW-VP7-R: GCTTCGTCTGTTTGCTGGTA

RIP-EW-NSP4-F: TGCACTGACTGTTCTATTTACGA

RIP-EW-NSP4-R: GGGAAGTTCGCATTGCTAGT

RIP-EW-NSP5/6-F: GGACACCGCAAGGTCAAAAA

RIP-EW-NSP5/6-R: TCGTCTGAGTCTGATTCTGCTT

### J2 Immunofluorescent staining

IECs from Mettl3^ΔIEC^ mice as well as from the wild-type littermate control mice were isolated as described in previous section. Isolated IEC were centrifuged onto glass slides and fixed with 4% Paraformaldehyde for 30 min at room temperature. Subsequently, permeabilized and blocked with PBS containing 0.1% Triton-X-100 and 5% bovine serum albumin for 1 h at room temperature. Double-stranded RNA (dsRNA) was labeled by a mouse monoclonal antibody J2 (Scisons) for 2 h at room temperature, followed by incubation with anti-mouse IgG Alexa Fluor 594-conjugated antibody (Invitrogen) for 1 h, and cells nuclei were visualized with 4,6-diamidino-2-phenylindole (DAPI, Invitrogen). All fluorescence images were analyzed via confocal imaging using Zeiss LSM880.

### Statistical analysis

Statistical analysis was performed with the GraphPad Prism 8.0 (GraphPad, Inc., USA). Experiments were independently repeated for indicated times listed in the figure legend. Representative data was exhibited as the means ± SEM. Quantitative data was compared using two-tail Student t test. In addition, correlational analysis of gene expression was conducted with linear regression. P-values for every result were labeled on figures, and P < 0.05 was reckoned as statistically significant (*P < 0.05, **P < 0.01, ***P < 0.001, ****P < 0.0001, NS., not significant).

## Data availability statement

The data that support the findings of this study are available from the corresponding author upon reasonable request. RNA sequencing data are available from the SRA database with accession numbers PRJNA713535.

## Author Contribution

A.W. designed, performed and interpreted experiments. W.T. analyzed the RNA sequencing data and m6A-Seq data. J.T. did the m6A-Seq experiment. J.W., J.G., G.Z., X.R., R.L., and D.W. helped with animal experiments and cellular experiments., S.D., R.A.F., K.Z., and W.P. provided critical comments and suggestions; A.W. and S.Z. wrote the manuscript. S.D. and W.T. edited the manuscript. S.Z. and H.B.L. supervised the project.

## Acknowledgements

We would like to thank Hongdi Ma, Taidou Hu, Kaixin He, Yinglei Wang, Ji Hu, Anlei Wang, and Meng Guo for technical help and helpful discussion.

---

**Figure s1.**
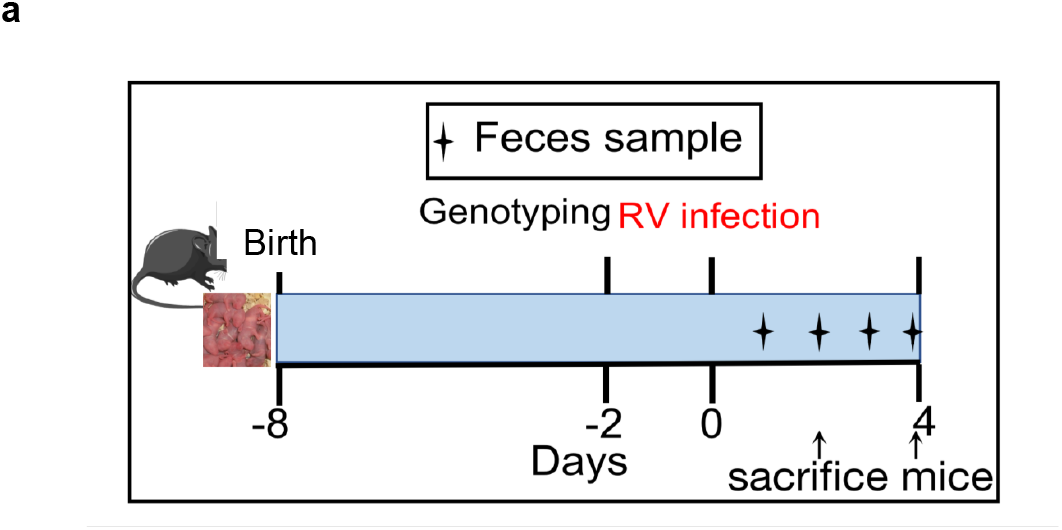
Schematic design of RV infection.

**Figure s2.**
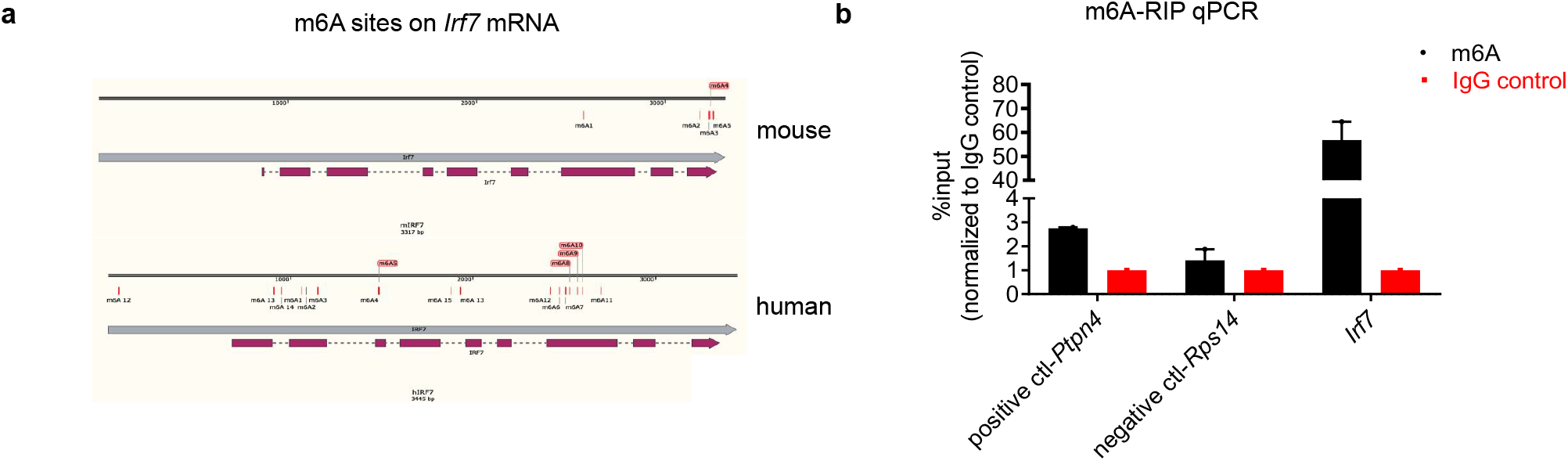
Characterization of m6A modifications on *Irf7* mRNA. (**a**) Predicted m6A sites of mouse or human *IRF7* on the genomes. (**b**) m6A-RIP-qPCR confirms *Irf7* as an m6A-modified gene in IECs. Mice were infected by RV for 2 days. Fragmented RNA was incubated with an anti-m6A antibody (Sigma Aldrich ABE572) or IgG IP Grade Rabbit polyclonal antibody(abcam, lot: 934197). The eluted RNA and input were processed as described in ‘RT-qPCR’ section, the data were normalized to the input samples (n=2, mean ± SEM), *Ptpn4* and *Rps14* were measured with m6A sites specific qPCR primer as positive control and negative control, Irf7 was measured with predicted m6A sites specific qPCR primer.

**Figure s3.**
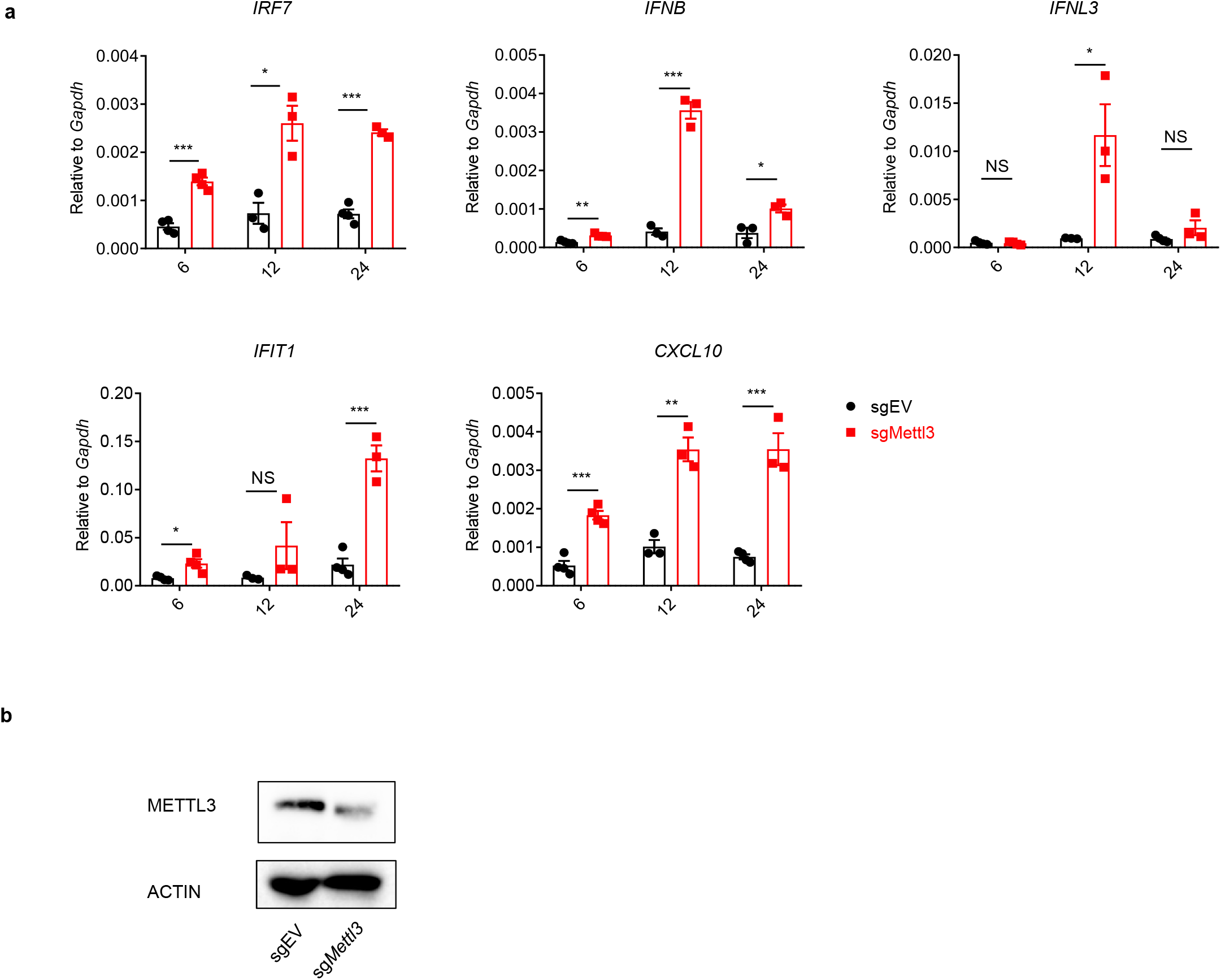
*METTL3* knockdown in HT-29 cells results in increased IFN response. (**a**) qPCR analysis of IFN/ISGs in Rhesus rotavirus-infected *Mettl3* WT and KD HT-29 cells at indicated hours post infection (hpi) (mean ± SEM), statistical significance was determined by Student’s t-test (*P < 0.05, **P < 0.005, ***P < 0.001, NS., not significant), at least three replicate experiments were performed. (**b**) Knock down efficiency of *METTL3* in HT-29 cells.

**Figure s4.**
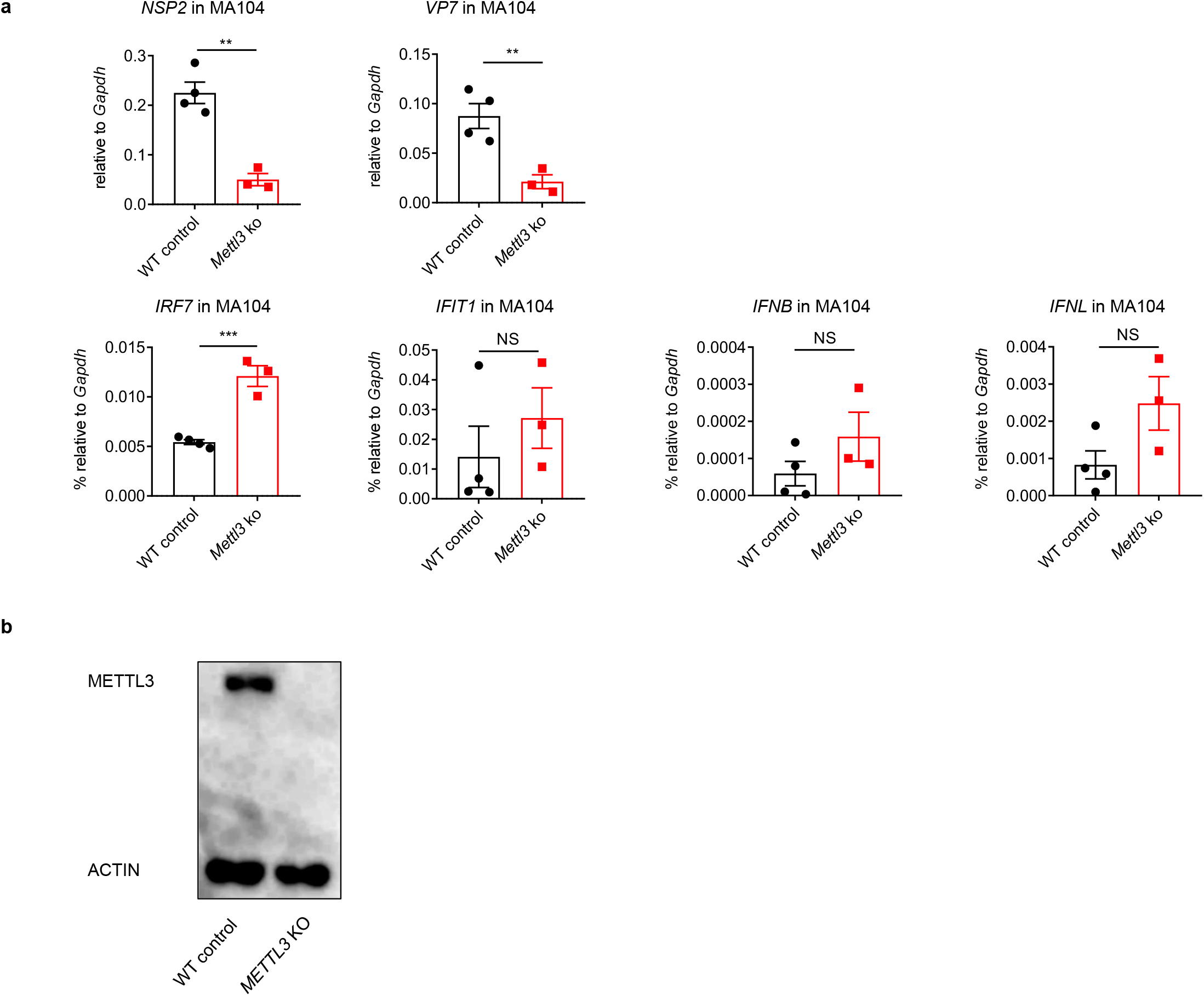
*METTL3* deficiency in MA104 cells results in increased resistance to Rhesus rotavirus infection. (**a**) qPCR analysis of viral RNAs, IFNs, and ISGs in Rhesus rotavirus-infected *Mettl3* WT and KO MA104 cells at 24 hours post infection (hpi) (WT control n=4, *Mettl3* ko n=3, mean ± SEM), statistical significance was determined by Student’s t-test (*P < 0.05, **P < 0.005, ***P < 0.001, NS., not significant). (**b**) Knock out efficiency of *METTL3* in MA104 cells.

**Figure s5.**
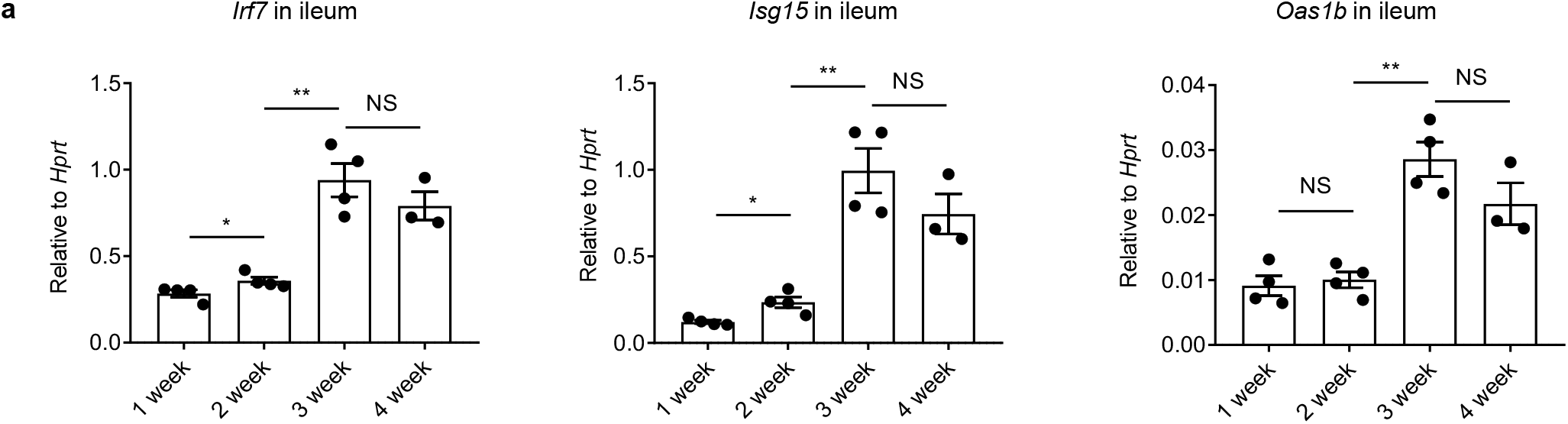
Expression of *Irf7* and *ISGs* in ileum from mice at the ages of 1-4 weeks post birth. (**a**) qPCR analysis of indicated genes in the ileum (1 week n=4, 2 week n=4, 3 week n=4, 4 week n=3, mean ± SEM). Statistical significance was determined by Student’s t-tests between ages (*P < 0.05, **P < 0.005, NS., not significant).

**Figure s6.**
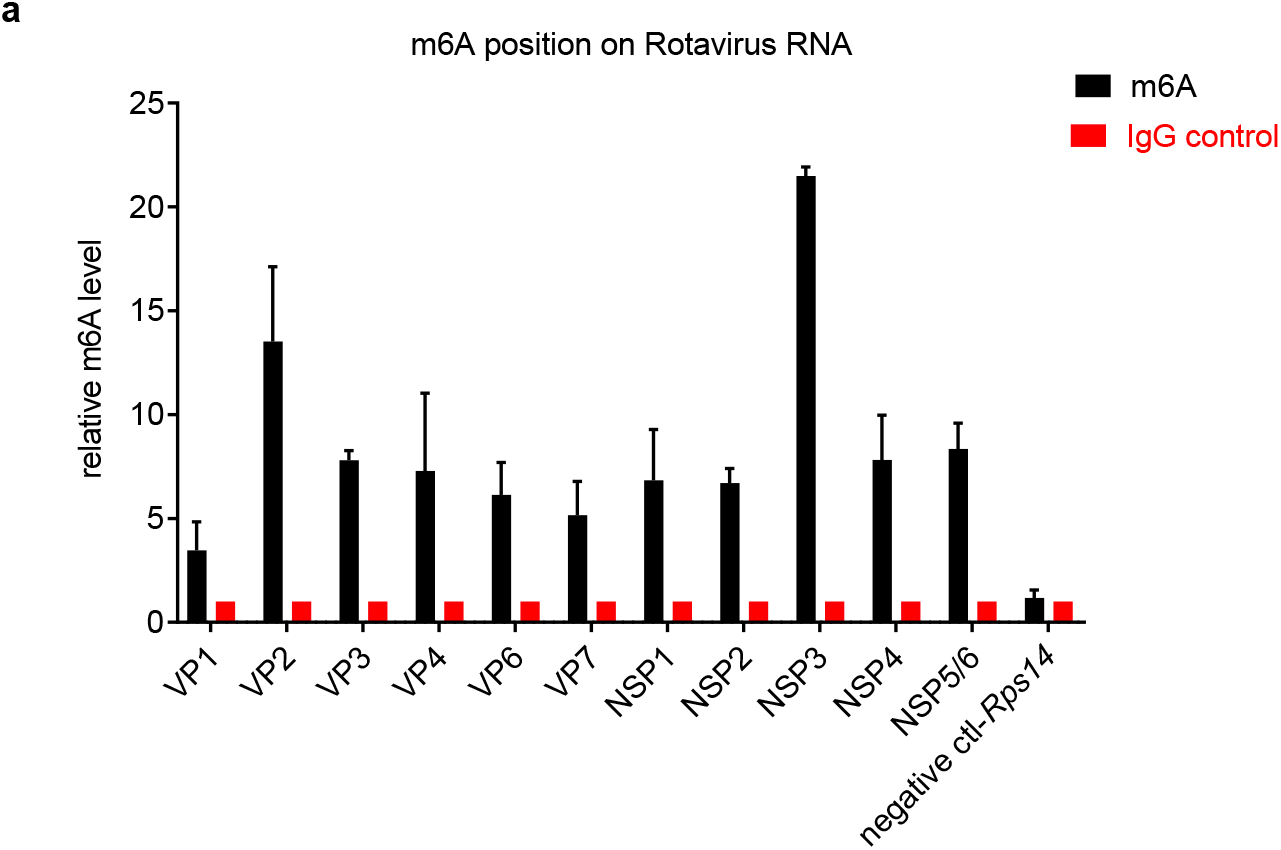
m6A-RIP-qPCR analysis of the predicted m6A sites on Rotavirus RNA. (**a**) Total RNA was extracted from IECs from 1-week-old WT mice infected by RV for 2 days. Fragmented RNA was incubated with an anti-m6A antibody (Sigma Aldrich ABE572) and IgG IP Grade Rabbit polyclonal antibody(abcam, lot: 934197). The eluted RNA and input were processed as described in ‘RT-qPCR’ section, the data were normalized to the input samples (n=2, mean ± SEM), Rps14 was chosen as a negative control.

**Figure s7.**
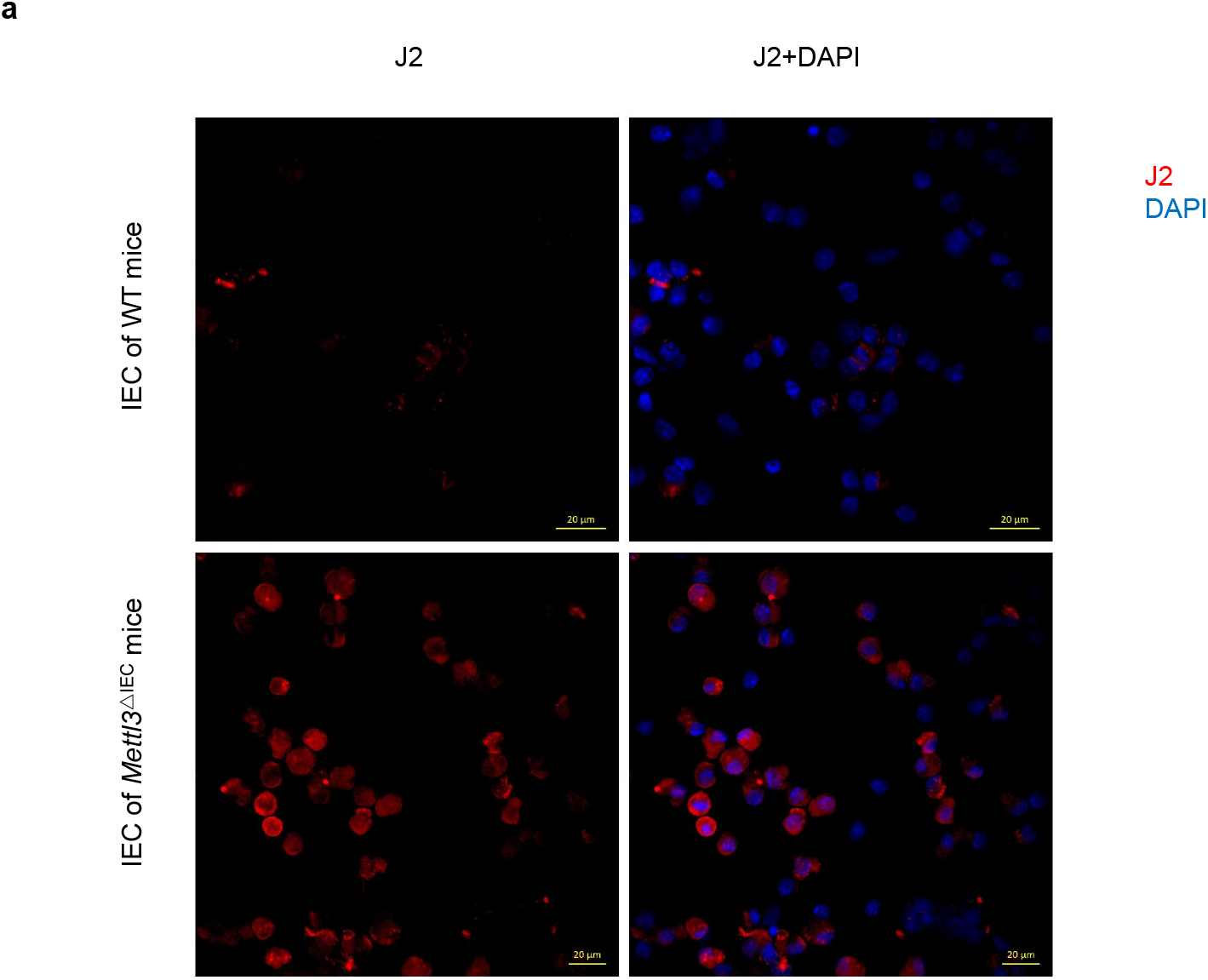
*Mettl3* deficiency leads to aberrant dsRNA formation in isolated IECs. (**a**) IECs from 6-weeks-old *Mettl3*ΔIEC mice as well as the WT littermate controls were isolated. Double-stranded RNA (dsRNA) was labeled by immunostaining with a mouse monoclonal antibody J2 (Scisons),cells nuclei were visualized with 4,6-diamidino-2-phenylindole (DAPI, Invitrogen). All fluorescence images were analyzed via confocal imaging using Zeiss LSM880.

